# HepT1-derived murine models of high-risk hepatoblastoma display vascular invasion, metastasis, and circulating tumor cells

**DOI:** 10.1101/2021.07.09.451809

**Authors:** Sarah E. Woodfield, Brandon J. Mistretta, Roma H. Patel, Aryana M. Ibarra, Kevin E. Fisher, Stephen F. Sarabia, Ilavarasi Gandhi, Jacquelyn Reuther, Zbigniew Starosolski, Andrew Badachhape, Aayushi P. Shah, Samuel R. Larson, Rohit K. Srivastava, Yan Shi, Richard S. Whitlock, Kimberly Holloway, Angshumoy Roy, Ketan B. Ghaghada, Dolores Lopez-Terrada, Preethi H. Gunaratne, Sanjeev A. Vasudevan

## Abstract

Hepatoblastoma (HB) is the most common pediatric primary liver malignancy, and survival for high-risk disease approaches 50%. Mouse models of HB fail to recapitulate hallmarks of high-risk disease. The aim of this work was to generate murine models that show high-risk features including multifocal tumors, vascular invasion, metastasis, and circulating tumor cells (CTCs). HepT1 cells were injected into the livers or tail veins of mice, and tumor growth was monitored with magnetic resonance and bioluminescent imaging. Blood was analyzed with fluorescence activated cell sorting to identify CTCs. Intra- and extra-hepatic tumor samples were harvested for immunohistochemistry and RNA and DNA sequencing. Cell lines were grown from tumor samples and profiled with RNA sequencing. With intrahepatic injection of HepT1 cells, 100% of animals grew liver tumors and showed vascular invasion, metastasis, and CTCs. Mutation profiling revealed genetic alterations in seven cancer-related genes, while transcriptomic analyses showed changes in gene expression with cells that invade vessels. Tail vein injection of HepT1 cells resulted in multifocal, metastatic disease. These unique models will facilitate further meaningful studies of high-risk HB.

**Summary Statement:** In this work, we developed and thoroughly characterized several unique models of hepatoblastoma derived from the HepT1 cell line that show high-risk features.

## Introduction

Hepatoblastoma (HB) is the most common pediatric primary liver tumor accounting for approximately 1% of all pediatric malignancies (Czauderna et al., 2014; Darbari et al., 2003; Hiyama, 2014). Notably, the incidence of HB has significantly increased in the past decade due to its association with premature birth (Spector and Birch, 2012), and this malignancy now has the fastest growing incidence of all pediatric solid tumors (Hubbard et al., 2019). HB generally afflicts children under the age of 4 years and has a 5-year overall survival (OS) rate of close to 80% for all patients (Czauderna et al., 2014). Unfortunately, high-risk patients that have multifocal, treatment refractory, metastatic, or relapse disease have a 5-year OS rate of only about 40% (Meyers et al., 2009). In general, HB is a genetically simple tumor with an average of only 2.9 mutations per tumor (Eichenmüller et al., 2014). The most commonly altered genes in HB are *CTNNB1* and *NFE2L2* including *CTNNB1* mutations or deletions in a majority of patients. Interestingly, mutations in *NFE2L2* are significantly correlated with metastasis, vascular invasion, and poor outcomes (Eichenmüller et al., 2014).

A challenge in the pre-clinical laboratory environment is developing cell line and murine models that depict high-risk characteristics of disease, such as vascular invasion, multifocal tumors, circulating tumor cells (CTCs), and metastasis, for meaningful studies. Our laboratory previously published two orthotopic xenograft models utilizing the established HepG2 and Huh-6 cell lines for intrahepatic growth of tumors, and these models recapitulate key features of disease, including generation of a robust blood supply in the tumor microenvironment, secretion of human α-fetoprotein (AFP), and immunohistochemical and transcriptomic similarity to primary tumors with expression of disease markers (Woodfield et al., 2017). At the same time, neither model showed obvious presence of high-risk features. Other standard models, including subcutaneous, splenic injection, genetically engineered (GEM), and hydrodynamic tail vein/Sleeping Beauty transposon models, also fail to consistently capture these high-risk attributes of disease (Whitlock et al., 2020). Use of patient-derived xenograft (PDX) models in the field is popular as it is thought that these models have fewer artifacts that arise with prolonged growth *in vitro* prior to implantation *in vivo* and instead faithfully recapitulate primary disease, including metastasis, but development of these models relies on a supply of fresh tissue samples from patients (Bissig-Choisat et al., 2016). To address this gap in disease modeling, we developed and thoroughly characterized several mouse models that are generated from the HepT1 cell line, which was previously established from a patient with HB (Pietsch et al., 1996) and is known to have a *CTNNB1* deletion and an *NFE2L2* mutation. Importantly, these models show key high-risk features of disease, including multifocality, CTCs, and metastasis, and will enable meaningful studies of these phenotypes that contribute to poor outcomes for HB patients.

## Results

### Intrahepatic injection of HepT1 cells generates invasive liver tumors

To generate HepT1-derived liver tumors, we injected two million HepT1 cells into the livers of immunodeficient NOD/Shi-*scid*/IL-2Rγ^null^ (NOG) mice (see methods). These cells were transduced with a lentiviral vector expressing the *luciferase* gene to allow longitudinal monitoring of tumors in living animals with bioluminescence imaging (BLI). Prior to implantation, we confirmed that the cells expressed strong luciferase activity (2-3 million relative luminescence (RLU)). Intraperitoneal injection of luciferin into the tumor-bearing animals caused the cells to emit a BLI signal that could be measured overtime (Figure 1A). Two of 9 animals (22%) emitted a BLI signal at 14 days after injection of cells, and 9 of 9 animals (100%) emitted a signal at 30 days after injection of cells (Figure 1A). Graph of the flux values shows their clear increase, indicating growth of tumors during the course of this experiment (Figure 1B). We also examined tumor burden in living animals with magnetic resonance imaging (MRI, Figure 1C) at early (4 weeks) and late (7 weeks) time points. We used contrast-enhanced MRI to examine angiogenesis of HepT1-derived tumors and found that they had grown a robust blood supply similar to tumors within humans (Figure 1D).

**Figure 1.**
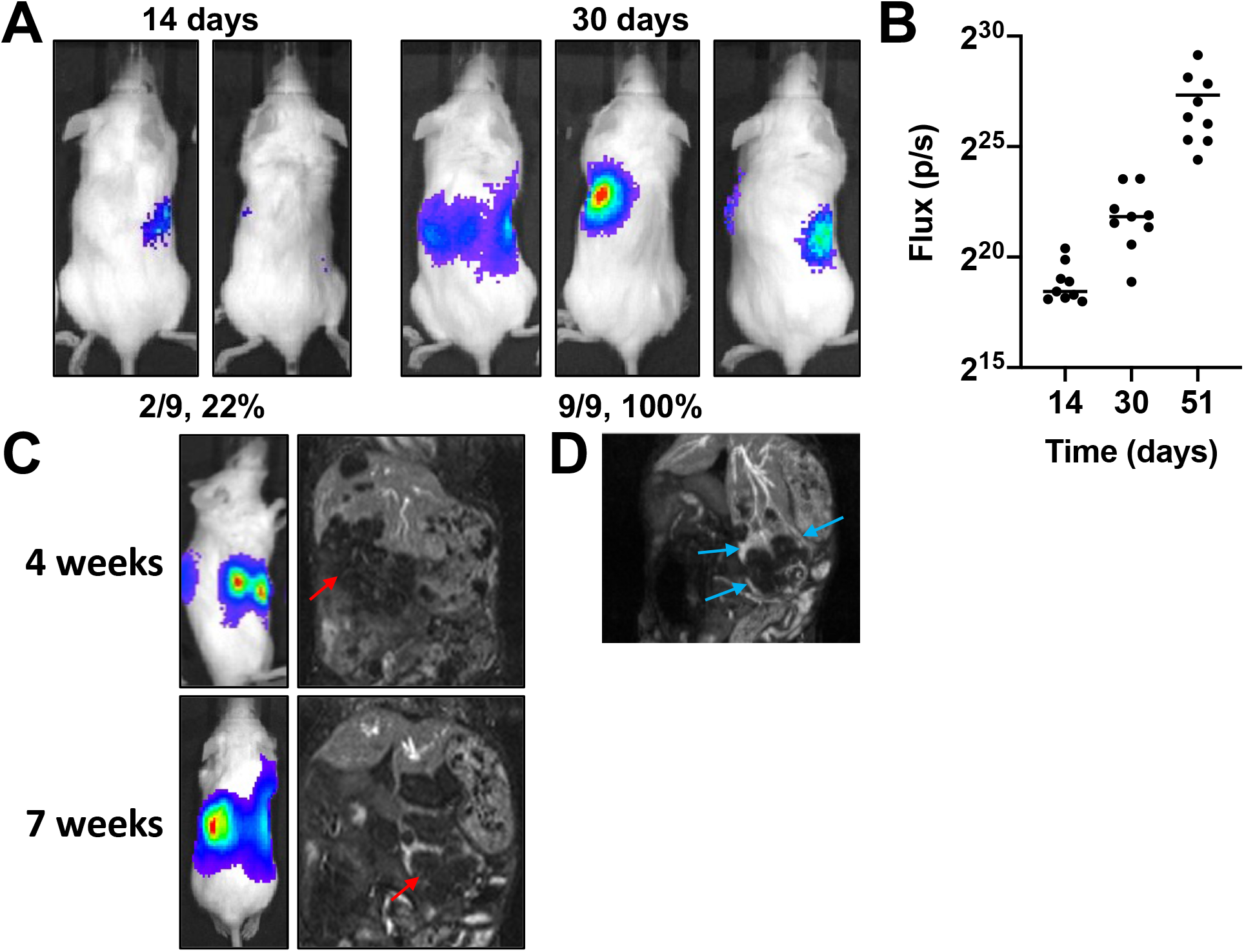
Growth of HepT1-derived tumors in living animals. (A) Nine animals were injected with HepT1 cells into the liver. At two weeks after injection, 2 of 9 (22%) of animals showed presence of tumor by BLI. At four weeks after injection, 9 of 9 (100%) of animals showed presence of tumor by BLI. Representative BLI images shown. (B) Increase in BLI signal shown at indicated time points after injection of cells. Flux (p/s) on y-axis shown with log2 scale. (C) Representative contrast-enhanced T1w-MRI images of animals harboring HepT1-derived tumors at early (4 weeks) and late (7 weeks) time points. (D) Contrast-enhanced T1-weighted coronal thick slab maximum intensity projection (MIP) abdominal images demonstrating the presence of increased vascularity (blue arrows) in the peripheral regions of a representative tumor.

After tumors were allowed to grow in animals for 7 weeks, the animals were euthanized and examined for liver tumor growth and also for presence of vascular invasion and extrahepatic disease. All animals that showed liver tumor with BL and MR imaging showed clear presence of intrahepatic tumors. Interestingly, we also found obvious presence of vascular invasion and extrahepatic disease within animals harboring HepT1-derived tumors. Shown in Figure 2A is a gross image of a liver tumor and attached areas of vascular invasion, including intrahepatic vascular invasion (IHVI) and inferior vena cava thrombus (IVCT). To further examine the tumor and areas of vascular invasion, we performed immunohistochemical staining of formalin-fixed paraffin-embedded (FFPE) tissues with hematoxylin and eosin (H&E) (Figure 2B). We also closely analyzed 8 animals for presence of specific extrahepatic disease at time of euthanasia and found that 6 of 8 (75%) of animals showed clear presence of extrahepatic disease, including peritoneal metastasis, pelvic metastasis, diaphragm metastasis, and abdominal wall metastasis. To look for lung metastasis, we serial sectioned FFPE lung tissues from these 8 animals and stained sections with H&E. We found obvious lung metastasis in one animal (Figure 2B). Interestingly, we were able to reimplant a whole piece of the IHVI sample into the liver of another immunocompromised animal to grow a primary liver tumor that resembled the primary tumors grown from injected cells. This was done utilizing a previously published technique used with patient-derived xenograft (PDX) tumors in which whole pieces are implanted with an incision in the Glisson’s capsule of the mouse liver and secured with a veterinary adhesive (Bissig-Choisat et al., 2016).

**Figure 2.**
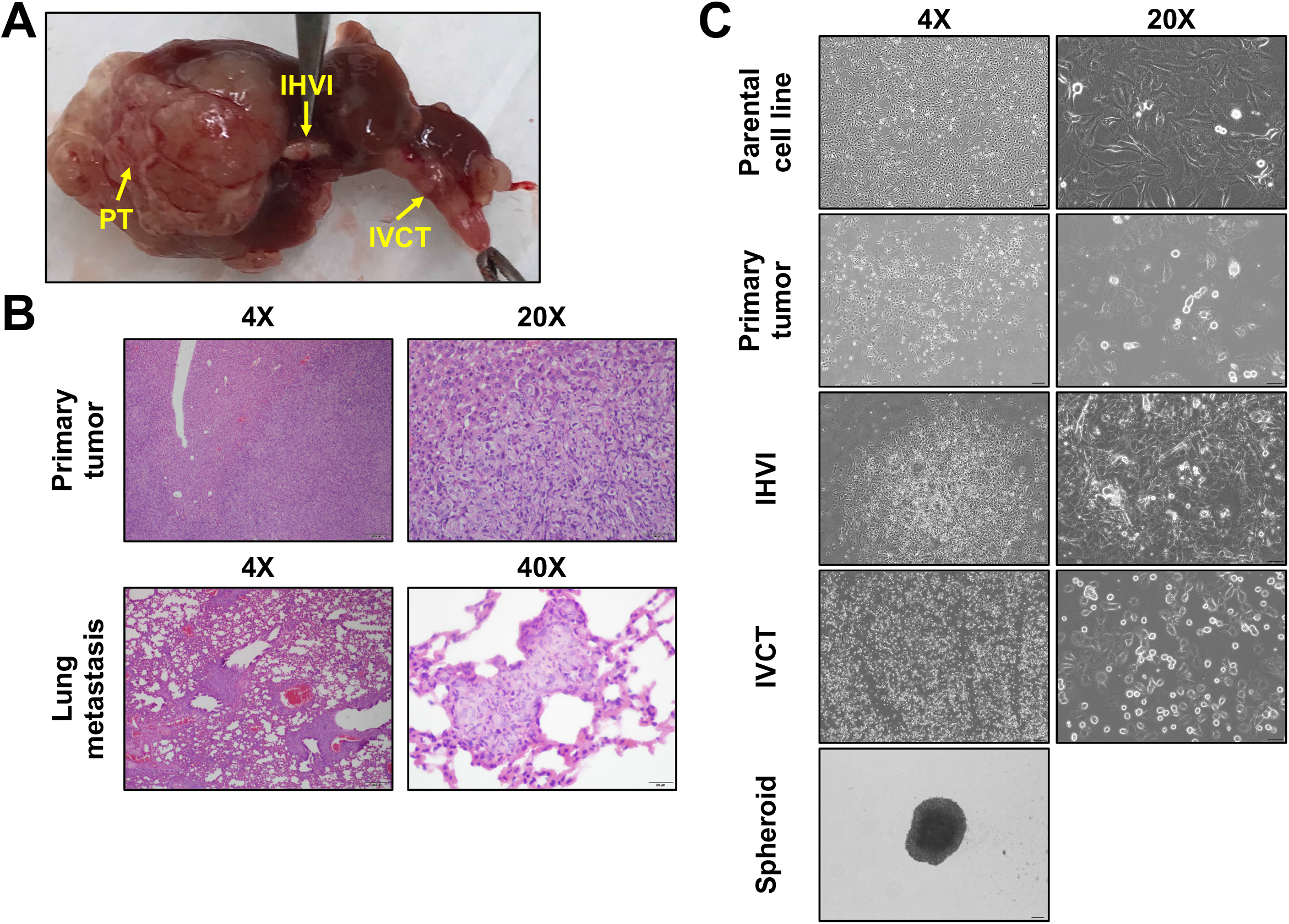
Intrahepatic injection of HepT1 cells leads to invasive disease. (A) Gross image of a HepT1-derived tumor with primary tumor (PT) and areas of vascular invasion (intrahepatic vascular invasion (IHVI), inferior vena cava thrombus (IVCT)) indicated. (B) Representative H&E images of primary tumor and lung metastasis. Scale bars represent 200 μm in 4X images, 50 μm in 20X images, and 20 μm in 40X images. (C) Brightfield images of the parental adherent cell line; cells grown from primary tumor, IHVI, and IVCT tissues in adherent conditions; and cells grown from primary tumor in spheroid conditions. Scale bars represent 200 μm in 4X images and 50 μm in 20X images.

Finally, with the primary tumor sample, the IHVI sample, and the IVCT sample, we grew cell lines in adherent and spheroid conditions. The primary tumor sample was grown in adherent and spheroid conditions to develop cell lines while the IHVI and IVCT samples generated two disparate adherent cell lines (Figure 2C). Interestingly, brightfield images of early passage adherent cells in culture show that these cell lines exhibit different visual phenotypes with the primary tumor cell line resembling the parental cell line and spreading across the plasticware while the IHVI- and IVCT-derived cells pile on top of each other as if they have lost their contact inhibition of growth (Figure 2C).

### Mutation profiling of HepT1 parental cell line

To verify the identity of the parental HepT1 cell line, we checked for the presence of known *CTNNB1* p.Ala5_Ala80del and *NFE2L2* p.Leu30Pro mutations (Eichenmüller et al., 2014). We ran a targeted next generation sequencing (NGS) custom pediatric cancer gene panel featuring 2247 coding exons of 124 genes as well as the *TERT* promoter. This work showed that our HepT1 cell line maintains the established *CTNNB1* and *NFE2L2* mutations (Figure 3, Table S1). Interestingly, this work also revealed mutations in four other genes with established clinical significance in pediatric solid tumors, *EP300, NF2, TP53*, and the *TERT* promoter, and loss of *PTEN* (Figure 3, Table S1). These mutations were confirmed with PCR and two-directional Sanger sequencing (Figure 3) with custom-designed primer sets (Table S2) (Sumazin et al., 2017).

**Figure 3.**
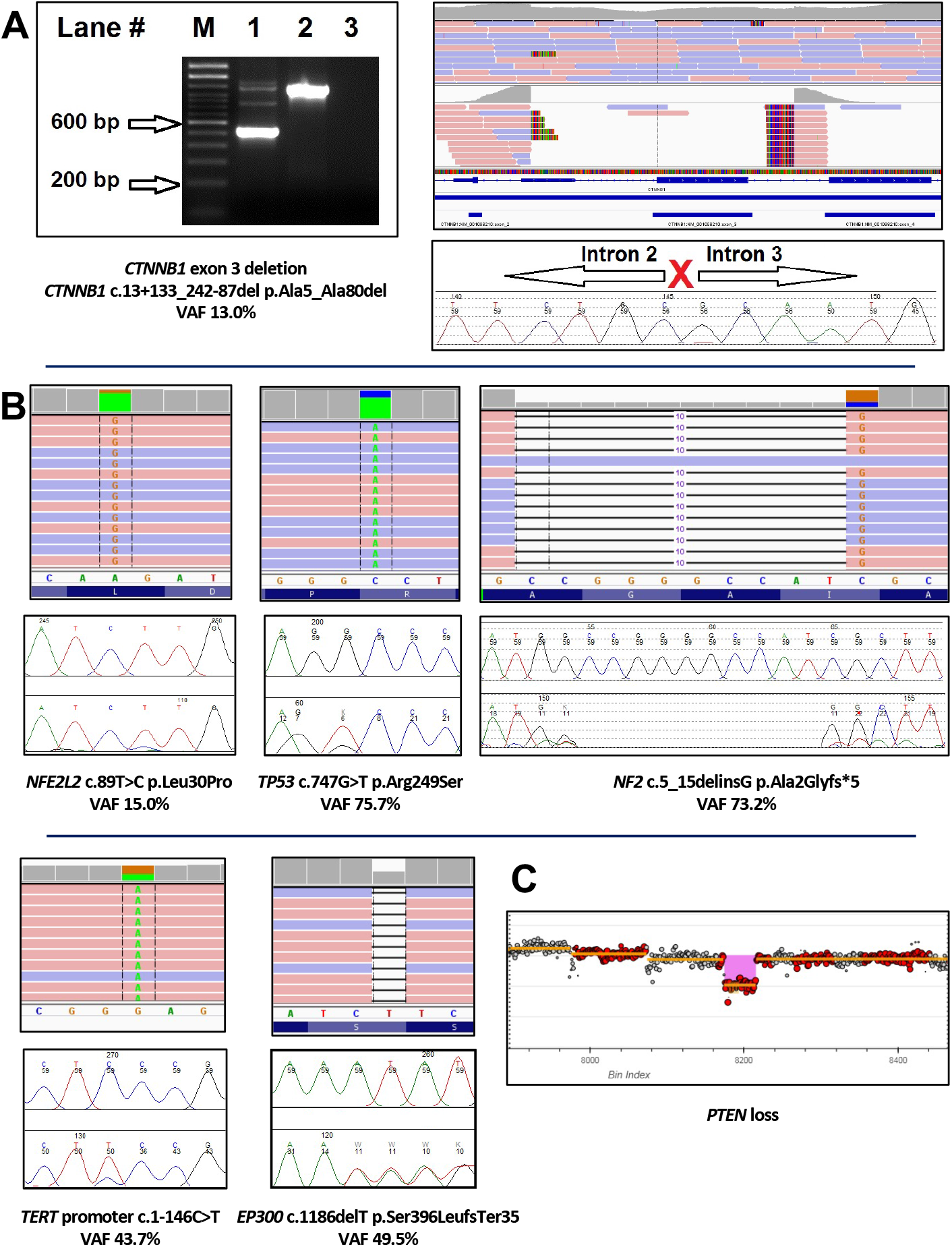
*CTNNB1 EP300, NF2, NFE2L2, TERT* promoter, and *TP53* mutations and *PTEN* loss detected in HepT1 cell line gDNA. (A) *CTNNB1* wild-type (1187 bp) and exon 3 deletion (533 bp) alleles are detected in HepT1 cells (Lane 1, 533 bp band) while only wild-type *CTNNB1* alleles are detected in A375 cells (Lane 2). An Integrative Genomics Viewer (IGV) version 2.4 representation of next-generation sequencing (NGS) pile-up up data shows the *CTNNB1* deletion beginning in intron 2, spanning exon 3, and ending in intron 3 which was confirmed by Sanger sequencing. (B) IGV 2.4 views of *NFE2L2, TP53, NF2, TERT* promoter, and *EP300* mutations detected by NGS and corresponding Sanger sequencing confirmations. (C) Copy-number changes consistent with *PTEN* loss were also observed although no orthogonal confirmation was performed. Gel Lane ID: #1 = HepT1 cells, #2 = A375 cells, #3 = No Template Control, M = 100 bp DNA ladder. Abbreviations: VAF, variant allele fraction.

### Transcriptomic profiling of tissue samples and cell lines grown from tissue samples

With this collection of eight unique HepT1-derived samples, including primary tumor (PT), IHVI, IVCT, IHVI grown as primary tumor, primary tumor adherent and spheroid cell lines, IHVI adherent cell line, and IVCT adherent cell line, we completed extensive total mRNA profiling with whole transcriptome next generation sequencing (RNA-seq). The principal component analysis (PCA) of this experiment shows three main clusters, one composed of the adherent cell lines, one containing only the IVCT sample, and one including the remaining tissue and spheroid cell line samples (Figure 4A). While the PC2 axis shows that the samples are similar, only showing differences in 19.2% of expressed genes among the samples, PC1 is more altered with differences in 31.7% of expressed genes between the IVCT sample and the other clusters. We speculate that this is due to a small amount of murine cell contamination (<10%) in the IVCT sample, which was removed from gene expression and pathway analyses moving forward. We also compiled this data in unsupervised hierarchical clustering heat maps to compare significant gene expression among the set of tissue samples and among the set of cell line samples (Figure 4B,C). Interestingly, the IHVI, IVCT, and primary tumor samples show clear differences in expression, and the IHVI grown as primary tumor sample more closely resembles the primary tumor sample than the IHVI sample (Figure 4B). This work also further shows that the three adherent cell line samples are much more similar in gene expression than the spheroid sample, which more closely resembles the primary tumor sample (Figure 4C).

**Figure 4.**
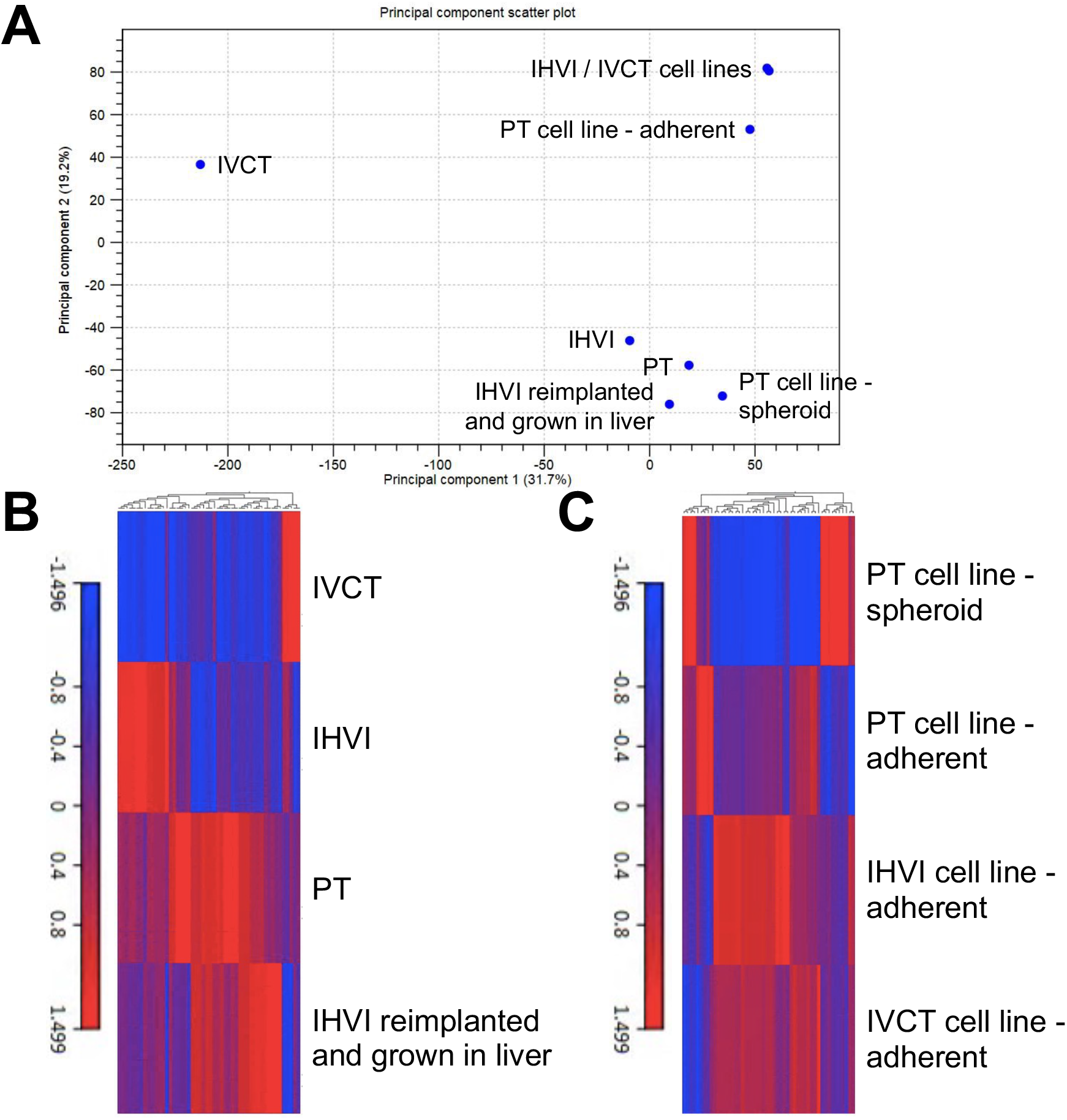
Transcriptomic profiling of HepT1-derived primary tumor and VI tumor sub-clones and cell lines. (A) PCA of all eight samples profiled, including tissue samples and cell line samples. X-axis represents PC1 and Y-axis represents PC2. (B) Hierarchical clustering heat map of differentially expressed genes with p-value < 0.05 from tissue samples. (C) Hierarchical clustering heat map of differentially expressed genes with p-value < 0.05 from cell line samples.

In a thorough assessment of differentially expressed genes, we found that only 44 genes show a consistent > 2-fold change or a < -2-fold change in each individual analysis of the IHVI, IVCT, and IHVI reimplanted samples compared to the primary tumor sample (Figure 5A). In contrast, the IVCT and IHVI cell line samples share 249 genes changed in common when compared to the primary tumor adherent cell line (Figure S1A). We then performed gene set enrichment analysis of the transcriptome data from the tissue and cell line samples using gene set enrichment analysis (GSEA) with the KEGG pathway enrichment database (Liao et al., 2019). GSEA on both IHVI samples compared to primary tumor and IVCT compared to primary tumor are shown in two separate bar graphs (Figure 5B). We see many similar upregulated pathways signifying a clear transcriptomic change from the primary tumor (Figure 5C). We also further examined the most significantly upregulated pathway among the vascular invasion tissue samples, the complement and coagulation cascades, in a hierarchical clustering heat map (Figure 5D). Interestingly, many of the genes in this KEGG pathway belong to two gene signatures used to molecularly characterize HB to predict prognosis (Cairo et al., 2008; Sumazin et al., 2017). We performed the same comparison of the adherent primary tumor cell line to the adherent IHVI and IVCT cell lines, respectively, and to the spheroid primary tumor cell line (Figure S1B-D). We see an overlap of 13 upregulated pathways, including the IL-17 signaling pathway that shows a > 2 enrichment score. Finally, we performed this same comparison of the adherent primary tumor cell line to the primary tumor and the spheroid primary tumor cell line to the primary tumor (Figure S2).

**Figure 5.**
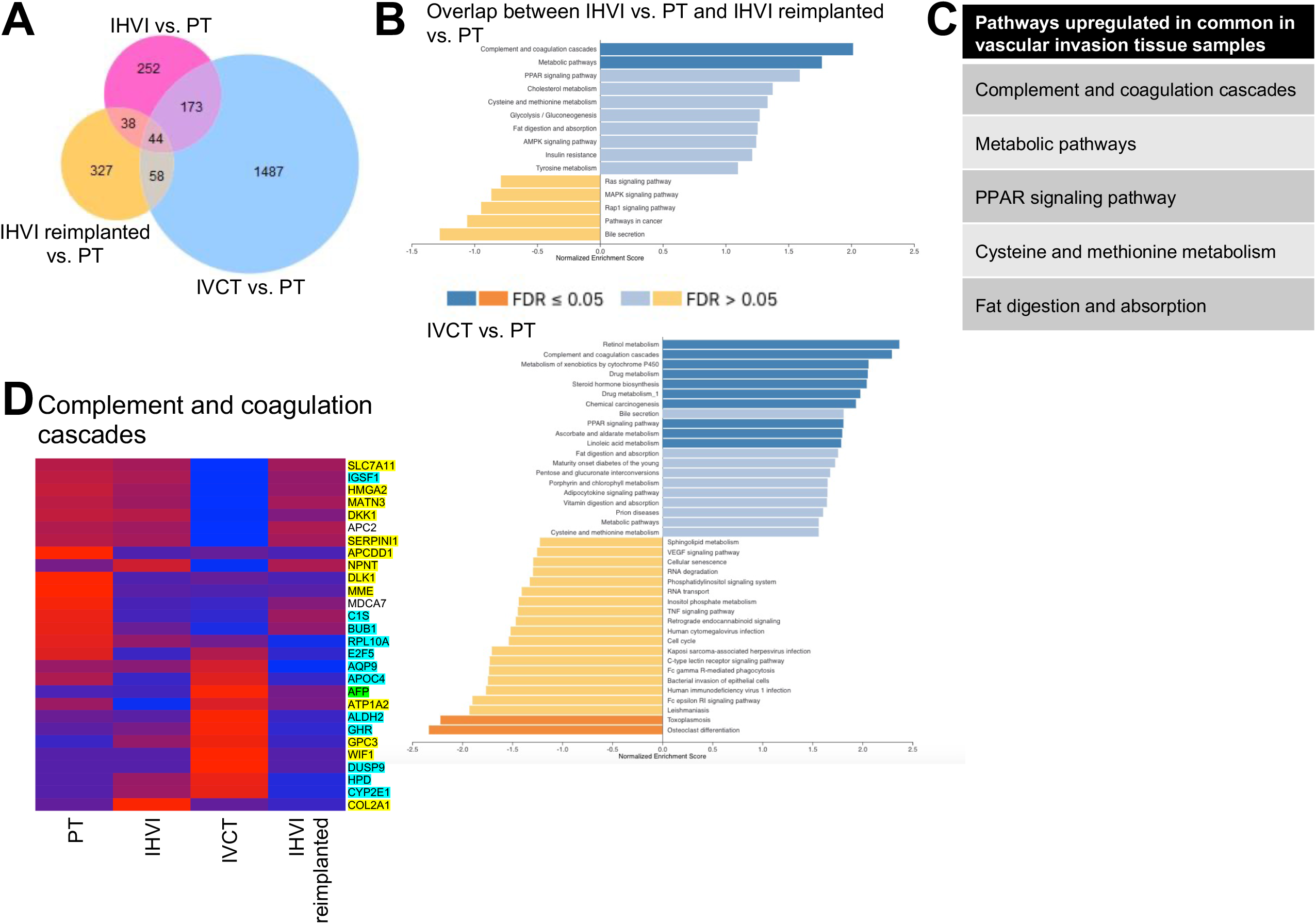
GSEA analysis of transcriptomic profiling. (A) Venn diagram of significantly changed genes with a p-value < 0.05 and log2 fold change > 1.5 or < -1.5 among the four tissue samples for input into KEGG pathway enrichment analysis. (B) Bar graphs of KEGG pathways significantly upregulated and downregulated in common between the primary tumor versus IHVI analysis and the IHVI grown as primary tumor versus IHVI analysis (top) and between the primary tumor and IVCT samples (bottom). (C) List of KEGG pathways upregulated in common among the vascular invasion tissue samples. (D) Heat map of the KEGG pathway most highly upregulated in the vascular invasion tissue samples, the complement and coagulation cascades. Highlighted genes are part of two gene signatures used for molecular classification of HB with those in yellow from Sumazin *et al*., 2017 (Sumazin et al., 2017); those in blue from Cairo *et al*., 2008 (Cairo et al., 2008); and those in green represented in both signatures.

### Detection of CTCs in the orthotopic HepT1 model

Because we saw clear presence of extrahepatic disease and lung metastasis in the original cohort of 9 animals, we undertook a second experiment with HepT1 cells transduced with a lentiviral vector expressing the *mCherry* gene for the red fluorescent protein to look for circulating tumor cells (CTCs). We injected two animals with two million cells into the liver to generate intrahepatic HepT1-derived tumors that express mCherry. After allowing the tumors to grow for 6 weeks, we collected blood from both tumor-bearing animals and also from four other unimplanted animals. We removed the plasma and red blood cells (RBCs) from the peripheral blood mononuclear cells (PBMCs) with standard density gradient centrifugation and RBC lysis and then analyzed these cells with fluorescence activated cell sorting (FACS) for mCherry to look for red fluorescent cells in the blood of the mice, which correspond to the CTC component. With both animals, we saw clear presence of mCherry-positive cells in the blood of animals (Figure 6A,D). We also imaged the animals at this time point with MRI and found that the raw number of mCherry-positive cells found in the blood correlated with the estimated tumor volumes by MRI (Figure 6C,D). Finally, at time of euthanasia, 10 weeks after implantation, we collected blood, separated the PBMC fraction, and plated the cells in culture to look for mCherry-positive cells. With fluorescence microscopy, we saw clear presence of mCherry positive cells from the blood of the animals (Figure 6B), which are HepT1-derived CTCs.

**Figure 6.**
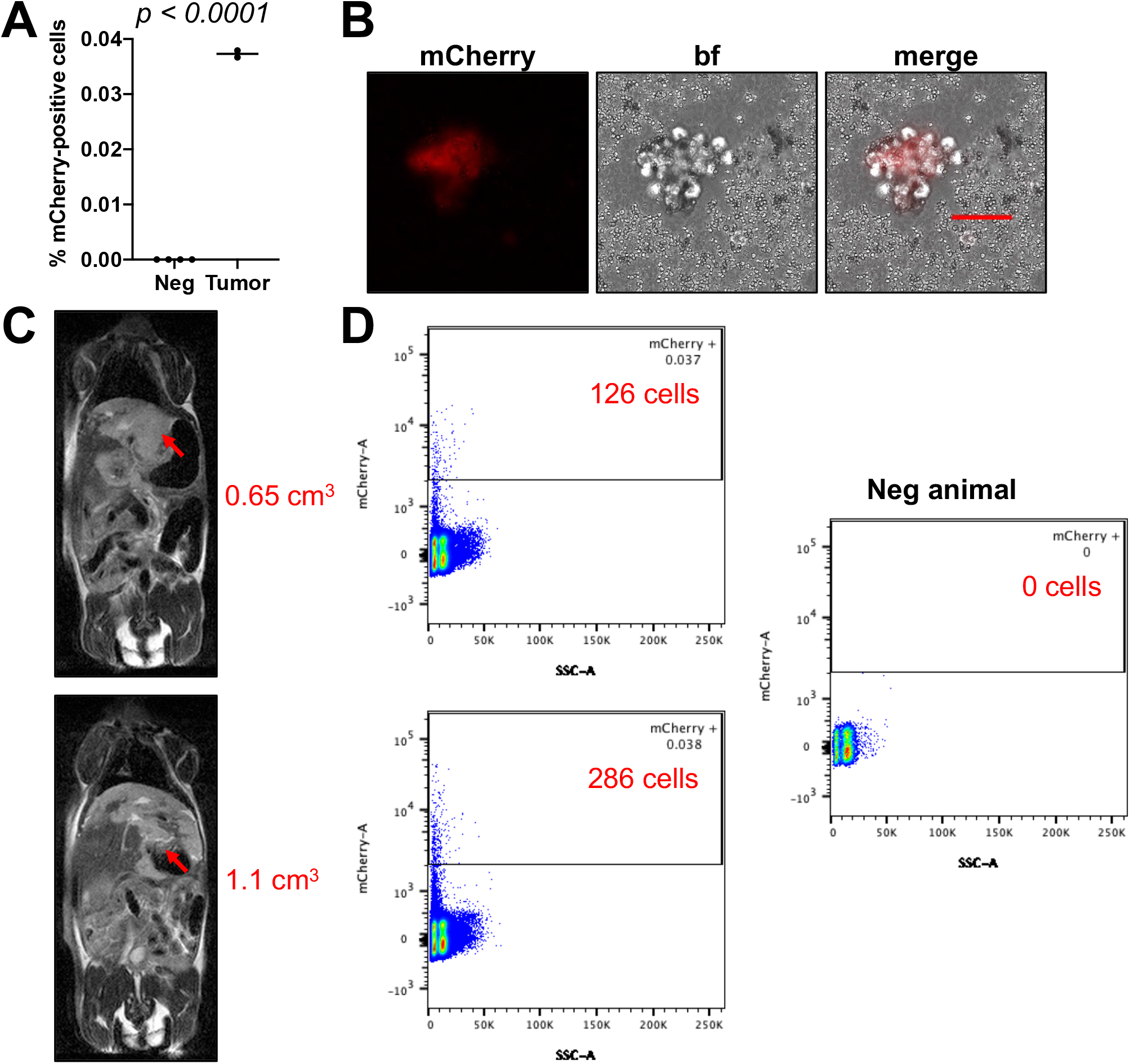
Detection of CTCs in the orthotopic HepT1 model. (A) Percent positive mCherry cells identified by FACS in blood of two mice harboring HepT1-derived tumors. Blood was taken 2 months after implantation of tumors. (B) mCherry-positive cells from the blood of one animal harboring a HepT1-mCherry-derived liver tumor. These cells were isolated at time of euthanasia 2 months after implantation of tumors. Scale bar represents 50 μm. (C) T2w-MRI images of two animals from which blood was taken at 2 months after implantation of tumors. (D) FACS plots graphed in A. Red labels indicate raw numbers of mCherry-positive CTCs detected. Plot of representative negative control animal not harboring a tumor indicated.

### Tail vein injection model of HepT1-derived liver tumors

In a final set of experiments, we generated a model of relapse HB with tail vein injection of HepT1-*mCherry* cells. In total, we injected six animals each with 1.5 million cells into the tail vein. Of these, 4 of 6 (66.67%) grew liver tumors, which we documented at time of euthanasia 12-13 weeks after implantation. Shown in Figure 7A is a representative gross image of a HepT1-derived tumor grown from tail vein injection of HepT1 cells, and shown in Figure 7D is a representative H&E image of such a tumor. These liver tumors are multifocal, as shown in the gross image of the tumor that has at least five independent tumor lesions (Figure 7A). We also looked for lung metastasis in the animals harboring HepT1-derived liver tumors and found an obvious lung nodule in a tumor bearing animal (Figure 7B). We checked this nodule with fluorescence microscopy and found that it was mCherry positive (Figure 7C), showing that it is derived from the HepT1-mCherry cells injected in the tail vein. With the animals harboring tumor, we collected the lungs for FFPE preservation, serial sectioning, and H&E staining. We found clear evidence of lung metastasis in all animals with a representative nodule shown in Figure 7D. Therefore, all animals that had liver tumors generated with tail vein injection also showed clear metastasis in the lungs. Taken together, this work shows that tail vein injection of HepT1 cells leads to repeated growth of multifocal liver tumors and lung metastasis that resembles relapse and metastatic HB seen in patients.

**Figure 7.**
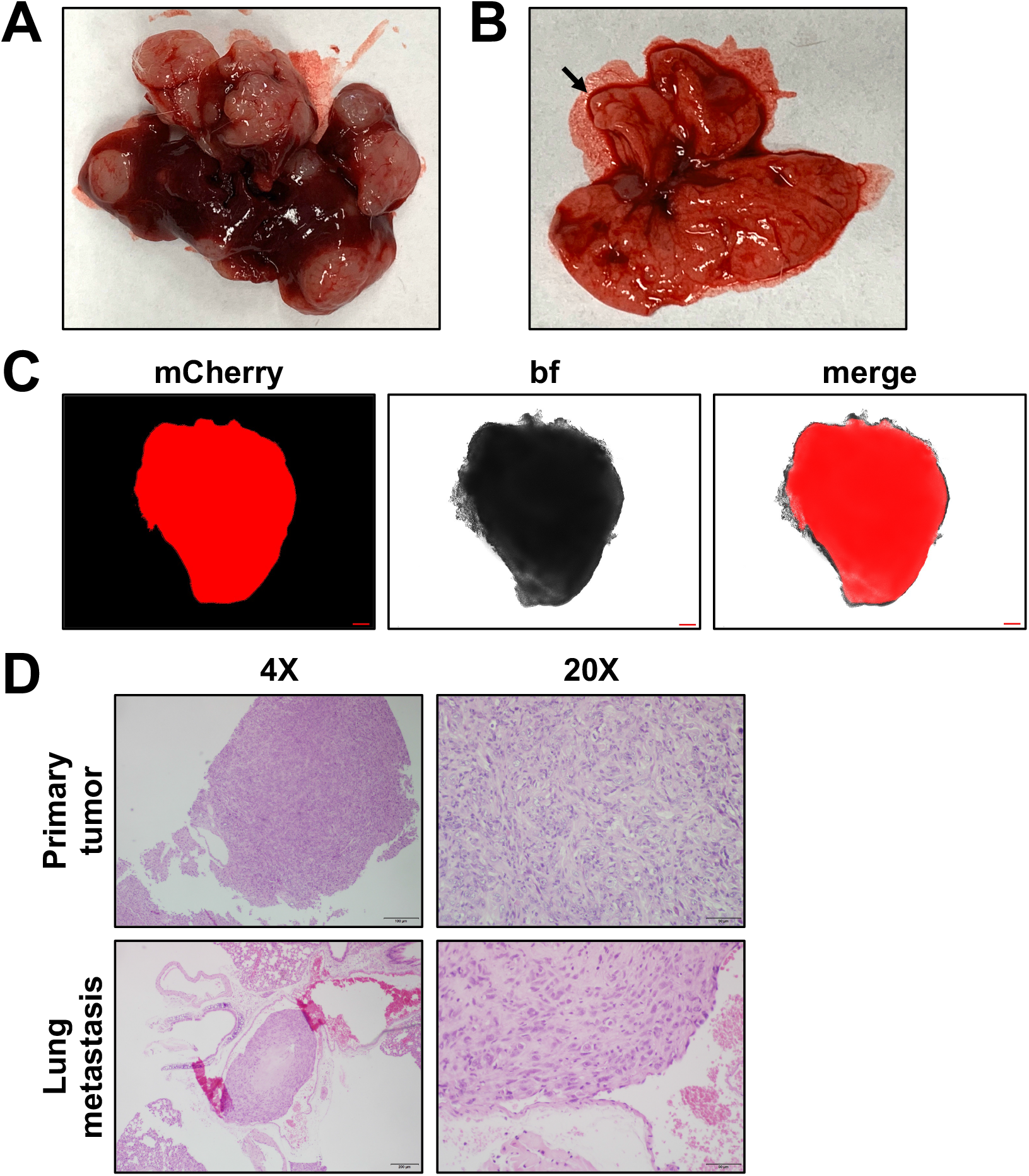
Tail vein injection HepT1-derived tumor model. (A) Gross image of a HepT1-derived intrahepatic tumor generated with tail vein injection of HepT1 cells. (B) Gross image of lung nodules from an animal tail vein injected with HepT1 cells. (C) Fluorescence image to show mCherry positivity of lung nodule shown in B. Scale bars represent 200 μm. (D) Representative H&E images of primary tumor and lung metastasis from animal harboring an intrahepatic tumor generated with tail vein injection. Scale bars represent 200 μm in 4X images and 50 μm in 20X images.

## Discussion

Overall, this work led to the development of three unique models of disease, including a *luciferase*-expressing intrahepatic injection model, an *mCherry*-expressing intrahepatic injection model, and an *mCherry*-expressing tail vein injection model. It is notable although not surprising that injection technique led to the generation of unique models that show different attributes of disease.

The intrahepatic injection model is usable for a wide range of studies, including, in particular, traditional drug studies that examine tumor growth in living animals and tumor size and phenotype after euthanasia. Notably, this model has an implantation rate of 100% 4 weeks after injection of cells, which is particularly helpful for studies in which replicate animals are required to show significant differences. In addition, this model gives rise to vascular invasion, metastasis, and CTCs, which are key hallmarks of high-risk disease. A recent retrospective review of vascular invasion in 66 patients with HB showed that vascular invasion is a prognostic indicator significantly associated with a worse 3-year OS and that patients without vascular invasion showed more favorable disease characteristics (Shi et al., 2017). It is also accepted in the field of cancer biology that a major complication and cause of cancer mortality is metastatic dissemination of primary tumor cells to distant sites by CTCs shed from the solid cancer tissue into the bloodstream of patients (Plaks et al., 2013). Thus, this model will allow particular examination of the mechanisms of these phenotypes in HB. Importantly this is the first report of HB-derived CTCs.

On the other hand, the tail vein injection model gives rise to disease that resembles relapse disease seen in patients with multifocal tumors spread throughout the liver and lung metastasis present in all animals that harbor primary liver tumors. Interestingly, in the literature, tail vein models are described as models of metastasis. When cells are injected through the tail vein, they first enter the venous circulation and confront the lung capillaries and then enter the arterial system and pass through the portal circulation. To our knowledge, most models generated with tail vein injection of cells lead to lung metastasis in the absence of primary tumors, and this modeling technique has been used with many solid tumor types including breast and lung cancers and melanoma (Gómez-Cuadrado et al., 2017). Thus, the tail vein injection model presented in this paper with HepT1 cells that leads to repeated growth of liver tumors and lung metastasis is very novel and shows a unique liver-specific tumor growth phenotype of this cell line.

Importantly, these models allow profiling of disease compartments, including primary tumor and areas of vascular invasion in this study but presumably also CTCs and metastasis in future work. This sequencing work is particularly devoid of background noise common to such “screening” experiments as all tissue and cell samples analyzed are derived from a common parental cell line. Interestingly, in the initial analysis of all eight samples, we saw clear separation of the IVCT sample. We speculate that this is due to contamination of the tumor thrombus sample with surrounding and embedded mouse blood cells. In any case, this represents < 10% of total analyzed mRNA and was removed for the additional gene expression and pathway analyses. Overall, deep analysis and validation of this data in future work will likely reveal clear drivers of vascular invasion and metastasis. The transcriptome sequencing analyses presented in this work already hint at a potential role for key pathways in vascular invasion in HB, including adaptive immune regulated responses, metabolic pathways, inflammatory cytokines, and Hippo signaling, which have already been shown to be involved in this process in other cancers. For example, mounting evidence shows that alterations in metabolism allow cancer cells to survive outside their solid tumor environments in blood vessels and facilitate them colonizing distant organs to generate metastasis (Elia et al., 2018). Furthermore, the upregulated retinol metabolism, drug metabolism, PPAR signaling pathway, cysteine metabolism, cytochrome P450 metabolism, and fat digestion and absorption pathways have been implicated in high-risk, metastatic HCC tumors, supporting the validity of these HepT1-derived models (Yi et al., 2019). Interestingly, levels of all but two of the genes identified to be significantly upregulated and downregulated that are part of the complement and coagulation cascades are also previously published to be predictive of patient prognosis as part of two molecular signatures of HB (Cairo et al., 2008; Sumazin et al., 2017). Finally, a specific analysis of transcriptomic data of HepT1 cells derived from the primary mouse liver tumors grown in adherent and spheroid conditions as surrogates for solid tumor primary tumor cells and floating CTCs revealed a potential role for IL-17 and TNF inflammatory cytokines in spheroid growth. Previous work has already implicated these cytokines in the support and promotion of breast cancer cell aggregate growth as spheroids (Geng et al., 2013). Overall, this work paves the way for potential future studies to search for therapeutic weaknesses of primary tumor cells, metastatic tumor cells, and floating CTCs. In studies of rare cancers, including pediatric solid tumors in general and especially HB, relevant laboratory models that accurately depict disease are even more necessary in the absence of large cohorts of patient data. These unique HepT1-derived models that exemplify high-risk disease features that contribute to poor survival rates will facilitate meaningful studies with the overall goal of improving patient outcomes.

## Materials and Methods

### Cells and culture conditions

The HepT1 cell line used in this study was obtained from Dr. Stefano Cairo. It was grown at 37°C in 5% CO_2_ in Eagle’s Minimum Essential Medium (EMEM, Lonza, Allendale, NJ) supplemented with 10% heat-inactivated fetal bovine serum (FBS, SAFC Biosciences, Lenexa, KS), 2 mM glutamine (Invitrogen, Carlsbad, CA), and 100 units/ml streptomycin/penicillin (Invitrogen). It was validated by DNA mutation analyses for known *CTNNB1* and *NFE2L2* mutations (see below). Cells that were harvested from tumors grown in mice or from mouse whole blood were maintained in adherent or spheroid plasticware in the same conditions as described above.

### Orthotopic mouse model with cells transduced with *luciferase* or *mCherry*

All animal procedures used in this study were performed under an animal protocol approved by the Institutional Care and Use Committee of Baylor College of Medicine (AN-6191). *In vivo* studies were performed in female 8 week old NOD/Shi-*scid*/IL-2Rγ^null^ (NOG) mice (Taconic Biosciences, Hudson, NY) similar to previous work (Woodfield et al., 2017). 2 × 10^6^ HepT1 cells transduced with *luciferase* or lentiviral psi-LVRU6MP-GFP-mCherry and resuspended in 25 μl phosphate-buffered saline (PBS) mixed with 25 μl matrigel (354230, Becton, Dickinson and Company) were injected into the right (*luciferase*) or left (*mCherry*) lobe of the liver through a right flank or midline abdominal incision, respectively. For the experiment with *luciferase*, the mice underwent BLI beginning at 10 days after implantation and every week thereafter with the *In Vivo* Imaging System (IVIS, PerkinElmer, Waltham, MA), and luminescence flux was recorded to assess tumor growth. After 7 weeks, necropsy was performed, intrahepatic and extrahepatic sites of tumor were noted, and samples were harvested for immunohistochemistry, RNA and DNA isolation, and to grow cell lines. For reimplantation of the intrahepatic vascular invasion area into the liver to grow a primary liver tumor, a previously published technique used with patient-derived xenograft (PDX) tumors in which whole pieces are implanted with an incision in the Glisson’s capsule of the mouse liver and secured with a veterinary adhesive (Bissig-Choisat et al., 2016). For the experiment with *mCherry*, 100 μl of blood was harvested from the facial veins of animals at 6 weeks after implantation for FACS to look for mCherry-positive CTCs. After 10 weeks, necropsy was performed, liver tumors and lung metastases were noted, and samples were harvested for immunohistochemistry. PBMCs from whole mouse blood were isolated with density gradient centrifugation and red blood cell (RBC) lysis (ACK lysing buffer, Gibco) and washed with cold 1X PBS. PBMCs were then analyzed with FACS or grown *in vitro* in non-adherent T-25 flasks in EMEM supplemented with 10% heat-inactivated FBS, 2 mM glutamine, and 100 units/ml streptomycin/penicillin as described above.

### Flow cytometry

PBMCs were analyzed with fluorescence activated cell sorting (FACS) on an Aria II (BD Biosciences) using a 355 nm UV laser to excite DAPI and a 450/50 nm bandpass filter to detect it and a 561 nm Yellow-Green laser to excite mCherry and a 610/20 bandpass filter to detect it. FSC and SSC were used to discriminate cell debris and 2 singlet discriminators were used to remove cells that adhered together. DAPI was used to extract dead cells. mCherry signal was measured after removal of cell clumps and dead cells. Student’s t-Test (two-tailed) was used to determine statistical significance.

### Tail vein injection mouse model with cells transduced with *mCherry*

All animal procedures used in this study were performed under an animal protocol approved by the Institutional Care and Use Committee of Baylor College of Medicine (AN-6191). *In vivo* studies were performed in male and female NOD/Shi-*scid*/IL-2Rγ^null^ (NOG) mice (Taconic Biosciences, Hudson, NY). 1.5 × 10^6^ HepT1 cells transduced with lentiviral psi-LVRU6MP-GFP-mCherry and resuspended in 100 μl phosphate-buffered saline (PBS) were injected into the tail vein. After 12 to 13 weeks, necropsy was performed, and liver and lung samples were imaged and harvested for immunohistochemistry.

### *In vivo* MRI

MRI was performed as described previously (Woodfield et al., 2017) on a 1.0 T permanent MRI scanner (M2 system, Aspect Technologies, Israel). A 35 mm volume coil was used for transmit and receive of radiofrequency (RF) signal. Mice were sedated using 3% isoflurane, setup on the MRI animal bed, and then maintained under anesthesia at 1-1.5% isoflurane delivered using a nose cone setup. Body temperature was maintained by circulating hot water through the MRI animal bed. Respiration rate was monitored using a pneumatically controlled pressure pad placed in the abdominal area underneath the animal. A long circulating liposomal-Gd blood pool contrast agent (SC-Gd liposomes) was systemically administered via the tail vein at a dose of 0.1 mmol Gd/kg and used for contrast-enhanced T1-weighted imaging (Ghaghada et al., 2017; Woodfield et al., 2017) shown in Figure 1. High-resolution contrast-enhanced MRI (CE-MRI) was performed using a T1-weighted 3D gradient echo (GRE) sequence with the following scan parameters: echo time (TE) = 3.5 ms, repetition time (TR) = 20 ms, flip angle = 70, slice thickness = 0.3 mm, field of view = 54 mm, number of slices = 68, matrix = 180 × 180, NEX = 1, in-plane resolution = 300 µm, scan time ∼5 min. Images were analyzed and processed in Osirix (version 5.8.5, 64-bit, Pixmeo, Bernex, Switzerland). Tumor volumes were segmented and 3D volume-rendered images were generated in Slicer (version 4.4.0) (Fedorov et al., 2012). For Figure 6, tumor imaging was performed using a T2-weighted fast-spin echo (FSE) sequence with the following scan parameters: echo time (TE) = 80 ms, repetition time (TR) = 4545 ms, slice thickness = 0.8 mm, field of view = 80 mm, number of slices = 24, matrix = 256 × 250, number of signal averages = 2, dwell time = 25 µs, scan time ∼4 minutes. Images were analyzed and processed in Osirix (version 5.8.5, 64-bit, Pixmeo, Bernex, Switzerland).

### H&E of tumor tissues

Tissue samples were fixed in 4% paraformaldehyde (PFA, Alfa Aesar, Ward Hill, MA) overnight at 4°C. Tissues were then dehydrated in 70% ethanol until processing in paraffin. Samples were processed for H&E in the Texas Medical Center Digestive Diseases Center (Houston, TX). Imaging of tumor sections on slides was done on a DMi8 microscope (Leica, Germany).

### Mutation analysis of cell lines and xenograft tumors

DNA and RNA was extracted from FFPE PDX tissue using the DNA FFPE Tissue Kit from (Qiagen) and RecoverAll Total Nucleic Acid Isolation kit (Ambion), respectively. For targeted DNA sequencing, 50ng of DNA was utilized to generate NGS libraries using the KAPA Biosystems HyperPlus kit, followed by hybridization capture using a custom-designed SeqCap Target Enrichment probe set (Roche) targeting coding and canonical splice regions (+/-2 bp) of 2247 exons in 124 genes and the *TERT* promoter. NGS libraries were pooled and sequenced on an Illumina MiSeq utilizing 600v3 chemistry with 150bp read lengths. FASTQ files were aligned to the GRCh37 (hg19) reference human genome using BWA v0.7.12 and NextGENe v2.4.1.2 with variant calling performed by NextGENe v2.4.1.2 and Platypus v0.8.1 for single nucleotide variants and small indels (<25 bp), and Pindel v0.2.5 and Delly v0.8.1 for longer indels (>25 bp). Variants were annotated with Variant Effect Predictor. Copy number variant (CNV) analysis was performed using CNVkit v.0.9.3 on BAM files with pooled normal as reference. Samples achieved on average 2.58 million unique paired-end reads with a mean coverage of 345x. Variant interpretation and classification was performed using established laboratory procedures. To confirm the mutations detected by next-generation sequencing (Table S1), PCR was performed with 125 ng of HepT1 gDNA using custom-designed primer sets (Table S2) (Sumazin et al., 2017). After 2.0% agarose gel electrophoresis to confirm the correct size of PCR products, two-directional Sanger sequencing was completed, and mutations were detected with Mutation Surveyor, version v.5.0.1 (Softgenetics, State College, PA), Results of the confirmations are provided in Figure 3.

### RNA sequencing of cell lines and tumor tissues

RNA from frozen cell lines was isolated using the Direct-zol RNA MiniPrep Kit (Zymo). RNA from frozen tissue samples was isolated using the RNeasy Plus Mini Kit (Qiagen). Samples were treated with DNase 1 and eluted in nuclease-free water. Extracted RNA samples underwent quality control assessment using the RNA tape on Tapestation 4200 (Agilent) and were quantified with Qubit Fluorometer (Thermo Fisher). RNA libraries were prepared and sequenced at the University of Houston Seq-N-Edit Core per standard protocols. RNA libraries were prepared with QIAseq Stranded Total RNA library kit (Qiagen) using 100 ng input RNA. Ribosomal RNA was depleted with QIAseq FastSelect HMR kit (Qiagen). RNA was fragmented, reverse transcribed into cDNA, and ligated with Illumina sequencing adaptors. Size selection for libraries was performed using SPRIselect beads (Beckman Coulter), and purity of the libraries was analyzed using the DNA 1000 tape Tapestation 4200 (Agilent). The prepared libraries were pooled and sequenced using NextSeq 500 (Illumina) generating ∼20 million 2 × 76 bp paired-end reads per sample.

### RNA sequencing transcriptome analysis

The RNA-seq raw fastq data was processed with CLC Genomics Workbench 12 (Qiagen). The Illumina sequencing adaptors were trimmed and reads were mapped to the HG38 human reference genome. Read alignment was represented as integer counts by using parameters of mismatch cost 2, insertion cost 3, length fraction 0.8, similarity fraction 0.8, max of 10 hits for a read. Integer read counts were normalized by Trimmed Means of M-values (TMM) algorithm. After normalization, we performed differential gene expression using the EdgeR package (Robinson and Oshlack, 2010) which uses a generalized linear model linked to the negative binomial distribution to identify significance. The significance level of FDR adjusted p-value of 0.05 and a log2 fold change greater than 1.5 or less than -1.5 was used to identify differentially expressed genes.

### GSEA of cell lines and tumor tissues

GSEA was performed using WebGestalt (web-based gene set analysis toolkit) for differentially expressed genes identified by EdgeR (Liao et al., 2019). The KEGG enrichment pathway database was used to profile the genes with a minimum of 5 genes per category and normalized enrichment to a significance level false discovery rate (FDR) < 0.05.

### Microscopy

Brightfield images of tumor sections were taken on a DMi8 microscope (Leica) at the indicated magnifications and scales. Brightfield images of cells and fluorescent microscope images were taken on a BZ-X710 All-in-One Fluorescence Microscope (Keyence, Itasca, IL, USA) at the indicated magnifications and scales.

## Acknowledgments

We would like to thank Dr. Stefano Cairo (XenTech, France) for the generous gift of the HepT1 cell line. We would like to thank Dr. Igor Stupin for assistance with MRI. We would like to thank Pamela Parsons in the Texas Medical Center Digestive Diseases Center for completing the immunohistochemistry experiments. This project was supported by the Cytometry and Cell Sorting Core at Baylor College of Medicine with funding from the CPRIT Core Facility Support Award (CPRIT-RP180672) and the NIH (CA125123 and RR024574) and with the expert assistance of Joel M. Sederstrom and Amanda N. White.

## Data Availability Statement

The data presented in this study are available on request from the corresponding author.

## Author Contributions

Conceptualization, S.E.W. and S.A.V.; methodology, S.E.W., R.H.P., and S.A.V.; validation, S.E.W., B.J.M., R.H.P., K.E.F., S.F.S., and S.A.V.; formal analysis, S.E.W., B.J.M., K.E.F., S.F.S., I.G., J.R., Z.S., A.B., K.B.G., D.L.T., P.H.G., and S.A.V.; investigation, S.E.W., B.J.M., R.H.P., A.M.I., S.F.S., I.G., J.R., Z.S., A.B., A.P.S., S.R.L., R.S., Y.S., R.S.W., and S.A.V.; resources, S.E.W., R.H.P., A.M.I., I.G., J.R., S.R.L., K.H., and S.A.V.; data curation, B.J.M., K.E.F., I.G., J.R., K.H., A.R., and P.H.G.; writing—original draft preparation, S.E.W., B.J.M., K.E.F., S.F.S., J.R., and A.P.S.; writing—review and editing, S.E.W., B.J.M., K.E.F., S.F.S., A.B., A.P.S., R.S., K.B.G., D.L.T., P.H.G., and S.A.V. ; visualization, S.E.W., B.J.M., S.F.S., Z.S., and A.B.; supervision, S.E.W., A.R., K.B.G., D.L.T., P.H.G., and S.A.V.; project administration, S.E.W., R.H.P., I.G., J.R., and K.H.; funding acquisition, S.E.W. and S.A.V. All authors have read and agreed to the published version of the manuscript.

---

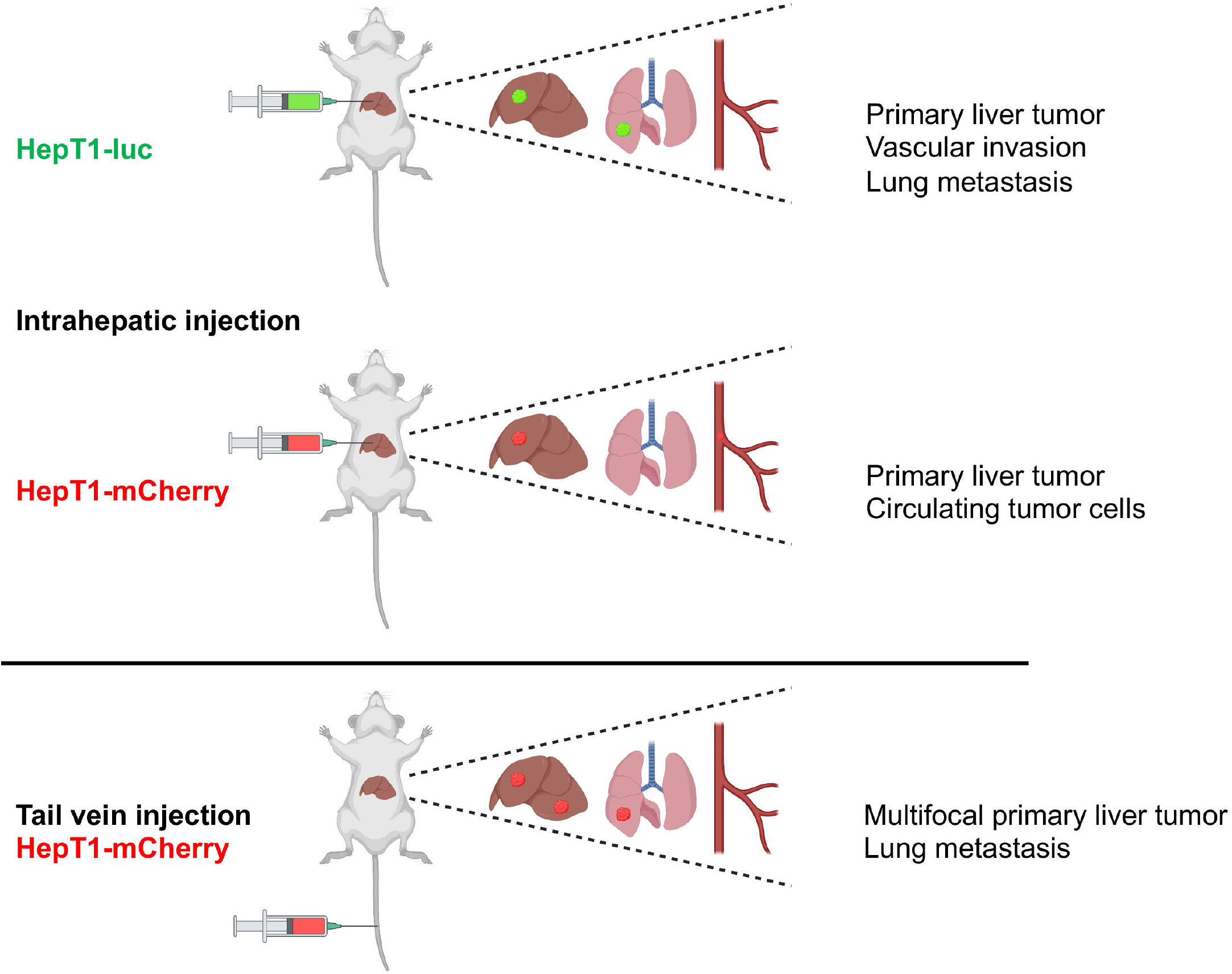

